# Metabolic profiling reveals nutrient preferences during carbon utilization in *Bacillus* species

**DOI:** 10.1101/2020.09.01.277376

**Authors:** James D. Chang, Ellen E. Vaughan, Carmen Gu Liu, Joseph W. Jelinski, Austen L. Terwilliger, Anthony W. Maresso

## Abstract

Pathogenic bacteria take host nutrients to support their growth, division, survival, and pathogenesis. The genus *Bacillus* includes species with diverse natural histories, including free-living nonpathogenic heterotrophs such as *B. subtilis* and host-dependent pathogens such as *B. anthracis* (the etiological agent of the disease anthrax) and *B. cereus*, a cause of food poisoning. Although highly similar genotypically, the ecological niches of these three species are mutually exclusive, which raises the untested hypothesis that their metabolism has speciated along a nutritional tract. Here, we employed a quantitative measurement of the number of reducing equivalents as a function of growth on hundreds of different sources of carbon to gauge the “culinary preferences” of three distinct *Bacillus* species, and related *Staphylococcus aureus*. We show that each species had widely varying metabolic ability to utilize diverse sources of carbon that correlated to their ecological niches. In addition, carbohydrates are shown to be the preferred sources of carbon when grown under ideal *in vitro* conditions. Rather unexpectedly, these metabolic utilizations did not correspond one-to-one with an increase in biomass, which brings to question what cellular activity should be considered productive when it comes to virulence. Finally, we applied this system to the growth and survival of *B. anthracis* in a blood-based environment and find that amino acids become the preferred source of energy while demonstrating the possibility of applying this approach to identifying xenobiotics or host compounds that can promote or interfere with bacterial metabolism during infection.

**Author summary:** Successful organisms must make nutritional adaptations to thrive in their environment. Bacterial pathogens are no exception, having evolved for survival inside their hosts. The host combats these pathogens by depriving them of potential biochemical resources, termed nutritional immunity. This places pathogens under pressure to utilize their resources efficiently and strategically, and their metabolism must in turn be tailored for this situation. In this study, we examined the carbon metabolism of three human pathogens of varying virulence (*Bacillus anthracis, Bacillus cereus*, and *Staphylococcus aureus*) and one nonpathogenic *Bacillus* (*Bacillus subtilis*) via a phenotype microarray that senses reducing equivalents produced during metabolism. Our analysis shows the existence of distinct preferences by these pathogens towards only a select few carbohydrates and implies reliance on specific metabolic pathways. These metabolic signatures obtained could be distinguished from one bacterial species to another, and we conclude that nutrient preferences offer a new perspective into investigating how pathogens can thrive during infection despite host-induced starvation.

## Introduction

One key hallmark of pathogens is their ability to use their hosts as a source of nutrients for survival and proliferation [1,2]. Bacterial pathogens, in term of their ecology, are bacteria that have undergone specialization to spend part or all of their lifecycle being dependent on their hosts for resources. This facilitates the use of host molecules for energy, catabolism/anabolism to build biomass, and replication of genetic material [3]. It is expected that bacterial pathogens adapt their metabolism to specifically exploit what the host offers; conversely, non-pathogenic bacteria could not exploit these resources but may better utilize nutrients in their abiotic environmental niche. Such fine tuning of metabolism would be advantageous, perhaps even essential, for pathogens to successfully carry out infection of the host. This competition between the host and pathogens for common resources offers insight into the functioning of nutritional immunity, a biochemical means of controlling bacterial pathogens that operate in conjunction with cellular immune systems [4-6].

*Bacillus anthracis* is the etiological agent of the deadly disease anthrax [7-9]. One of its more defining features is its ability to replicate to very high numbers in mammalian blood and tissues. As such, *B. anthracis* is often used as a model bacterial pathogen for the study of host nutrient uptake during infection [10-12]. Its infectious cycle begins when spores enter the host through an open wound, is inhaled, or is ingested. Next, spores germinate inside the host into the fully-replicative and growing vegetative cells. This life cycle is in stark contrast to *Bacillus cereus*, another member of the *Bacillus* genus, which is 93 percent similar at the genomic level to *B. anthracis* but known more for being a cause of food poisoning [13-15]. Another extensively studied *Bacillus* species, *Bacillus subtilis*, a non-pathogenic soil-dwelling bacteria that is utilized for food fermentation and as a biotechnology model system, is phylogenetically distinct from pathogenic *Bacillus*, as evidenced by sharing less than 20 percent of the amplified fragment length polymorphism markers, nor does it have any genes that code for known virulence factors [16-18].

Most of species in the genus *Bacillus* live ubiquitously in the environment similar to *B. subtilis*, and all except two of them (*B. anthracis* and *B. cereus*) are nonpathogenic to mammals. The extreme pathogenicity and virulence of *B. anthracis* is particularly striking when compared to other *Bacillus* species. It is largely believed that two additional genomic elements, the plasmids pXO1 and pXO2, which are not observed in other *Bacillus* species, are responsible for the virulence of *B. anthracis* [19,20]. In fact, transformation of these virulence plasmids into certain biovars of *B. cereus* has been demonstrated to result in bacteria that can cause anthrax-like disease [21,22]. Indeed, these plasmids encode for anthrax toxin and the poly-D-glutamic acid capsule, both of which are considered important virulence factors for the induction of anthrax, while pXO1 also codes for the transcriptional regulator *AtxA* which is known to control the production of toxin and S-layer [23-25]. However, most of the research into these plasmids thus far have been focused on production of toxins and capsule, and their effects on other aspects of *B. anthracis* biology, especially metabolism, remain undercharacterized. Given that the production of toxins must involve the survival and proliferation of the pathogen, we must also consider metabolism that fuels bacteria as being an essential part of virulence.

One approach to examining the role of metabolism in pathogenesis would be measuring the utilization of various nutrients by bacteria, ideally under conditions that mimic the host environment. This leads to the question of how nutrient utilization differs between pathogens and their nonpathogenic counterparts, especially in the genus *Bacillus*. Previous investigations of *B. anthracis* metabolism in association with virulence have thus far focused on roles of individual enzymes, a global genomic analysis, or characterization of metabolic regulators [26-31]. Here, we took a more comprehensive approach and assessed 189 distinct sources of carbon for their ability to drive the generation of reducing equivalents (here a proxy for metabolic outflow) for three species of *Bacillus* (*B. anthracis, B. cereus*, and *B. subtilis*) and *Staphylococcus aureus*. A pan-cupboard of optimal but also detrimental nutrients are reported that can be used to both enhance and reduce virulence and highlight how metabolism is specifically tailored along environmental niches.

## Results

### Quantification of bacterial carbon utilization through a colorimetric assay

Metabolic activity is powered by the breakdown of biologically useful molecules via the conversion of chemical potential energy into reducing potential energy [32]. For chemoheterotrophic bacteria that rely on carbon molecules as nutrients, one product of metabolism is the reductant NADPH. The quantity of intracellular NADPH can be measured colorimetrically through reduction of tetrazolium dyes that impart purple color [33-39]. The level of color is thought to be proportional to the overall metabolic activity, especially in terms of the generation of reductive potential. We hypothesized that the formation of NADPH in the presence of exogenously supplied nutrients might reflect different nutritional preferences between pathogenic and non-pathogenic bacteria. In this context, we assessed the metabolism of 189 different carbon sources for four different species of bacteria; *B. anthracis, B. subtilis, B. cereus*, and *S. aureus*. The experimental design of this study is shown in Fig 1A. We first aimed to determine some of the quantifiable parameters of the system, including the kinetics and endpoints of metabolism. Shown in Fig 1B is a plot of the metabolic activity (as measured by the reduction of tetrazolium) against time for three nutrients that display some of the types of activity curves observed in the data set. The first type of curve, shown here with D-glucose, is one in which the maximum rate of metabolic activity is observed for most of the experiment (green line). The second type, observed here with L-proline (pink line), shows classic exponential kinetics with an accelerated rate of metabolism followed by a slow saturation. Finally, many metabolites either inhibit metabolism or do not stimulate it, with data that resembles the curve shown for 2-hydroxy benzoic acid (blue line). In the analysis, we focused on two characteristic descriptors of metabolic activity: the metabolic endpoint, which represents the net colorimetric change over the course of the experiment, and maximum metabolic rate, which represents the highest rate of colorimetric change at all times. To average out background variation in the colorimetric measurement, the exponential moving average was employed to calculate a value for the final metabolic endpoint. To determine the maximum metabolic rate from a metabolic activity curve with multiple inflection points and stochastic variations, a polynomial was first fitted to the metabolic curve, and the resulting polynomial differentiated to give rate of metabolic change for all time points (see Materials and methods). We employed the use of two ready-made, commercially available plates with different sources of carbon (S1 Table) [33]. In this backdrop, all other nutrients in the system are not prominent sources of carbon. This was performed for 189 nutrients for three different *Bacillus* species (*B. anthracis, B. cereus*, and *B. subtilis*) as well as the related Gram-positive pathogen *Staphylococcus aureus*. There were striking differences in the both the maximum metabolic rate and maximum metabolic endpoint values between each species and each temperature (Fig 1C). Interestingly, whereas *B. anthracis* showed enhanced metabolism at the higher of the two temperatures (mean maximum metabolic rate for all nutrients at 30° = 10.93, at 37° = 27.19, p < 0.0001, paired Student’s t-test), *B. cereus* showed enhanced metabolism at the lower of the two (mean maximum metabolic rate for all nutrients at 30° = 16.77, at 37° = 6.52, p < 0.0001, paired Student’s t-test), a finding that may reflects adaptation of *B. cereus* for limited growth, multiplication, and sporulation in soil at lower temperatures (however, unlike true soil microorganisms it is not well adapted for using chemical resources, and is dependent on decaying organic matters for resources) [13]. The metabolism of *S. aureus* at body temperature was more similar to the metabolism of *B. anthracis* at body temperature than it was to *B. cereus*, presumably reflecting the ability of these organisms to infect a wide range of vertebrate hosts, and at different bodily sites (S1 Fig). We did not assess *B. subtilis* at higher temperatures because of its poor growth at 37 degrees (data not shown).

**Fig 1.**
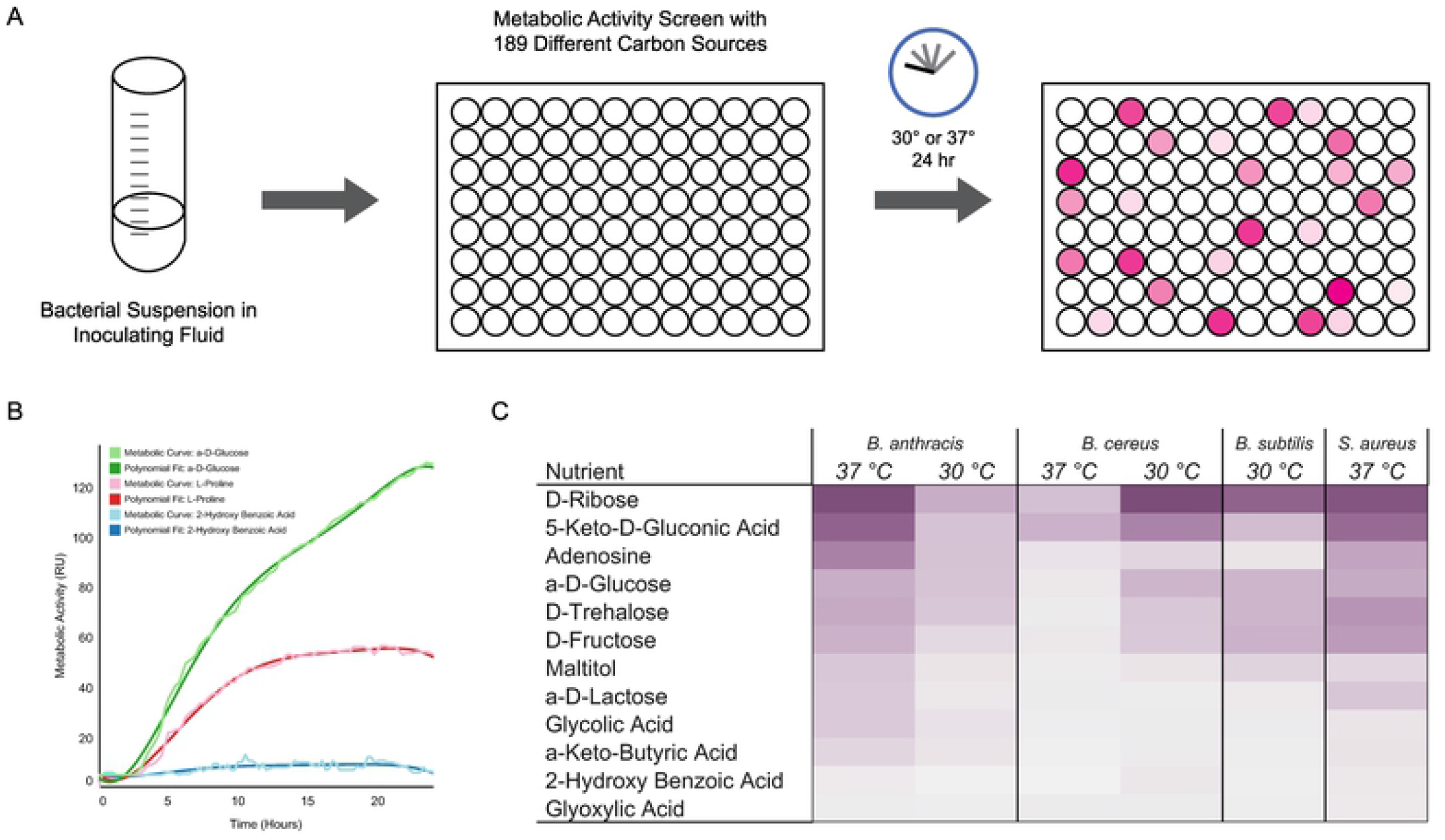
Colorimetric assay reflects metabolic activity in bacteria. (A) Schematic showing the experimental setup using 96-well plates with nutrients providing a carbon source for bacteria being examined. (B) Examples of raw metabolic data outputs and polynomial fitting for metabolic curves. Metabolic curves over the course of experiment for three nutrients with different degrees of color change are shown: High activity (green) with a-D-glucose, medium activity (red) with L-proline, and low activity (blue) with 2-hydroxy benzoic acid. Light curves show raw metabolic data output as measured by the overall color change, and corresponding dark curves show polynomials fitted to determine metabolic rates. (C) Maximum metabolic rates of bacteria and conditions tested for selected nutrients. Maximum metabolic rates for twelve selected nutrients from the carbon utilization screen are shown to highlight the range of rates measured. Darker shades reflect higher rates, and lighter shades lower rates. Two experiments in separate temperatures (30°C and 37°C) were performed for *B. anthracis* and *B. cereus* and are shown in two columns. Maximum metabolic rates are averaged from three independent runs.

### Bacterial metabolic activity and correlation to growth

Increases of the optical density at 600 nm in culture is typically used as a proxy for bacterial growth. We wished to also understand the relationship between bacterial growth and metabolism for nutrients assessed in Fig 1 across all three bacillus species. Rather remarkably, there was very little over-all correlation between optical density and metabolism for all compounds tested (S2 Fig). Spearman’s rank correlation coefficient (Spearman’s ρ) was calculated between rank ordered lists to ascertain the degree of correlation between these two metrics for metabolism. *B. anthracis-*ranked lists had the lowest correlation with ρ = 0.4753, while *B. cereus* and *B. subtilis* showed more similarity with ρ = 0.4810 and 0.5913 respectively. These values indicate that total metabolic activity as measured by chemical reductive potential does in some cases reflect enhanced growth of the organism, but in many other cases, it does not. Indeed, there were cases whereby very little increase in growth was observed (5-keto-D-gluconic acid) but reductive metabolism was one of the highest of all compounds tested (see *B. anthracis*) and other cases whereby growth was high (*B. subtilis* in capric acid) but almost no reductive metabolism was detected. Furthermore, these trends were not conserved amongst each species (despite strong reproducibility within each species), indicating that bacteria in *Bacillus* have vastly different species-specific metabolic programs that can run independent of its drive to replicate.

### Overall trends in metabolic utilization of carbon sources

We sought to determine whether the maximum metabolic rate could be used as a metric to compare different bacteria and under different conditions. Metabolic data were first standardized by each bacterium and condition to a mean of 0 and standard deviation of 1, and the resulting data were hierarchically clustered for organization. Data for bacteria incubated at their optimal temperature were used when two different temperatures were tested. When visualized as heat maps, metabolic rates showed that while few nutrients were well utilized in all bacteria, there also exists a group of nutrients that were utilized exceptionally by one species alone while not being used for metabolism in another species (Fig 2Ai and 2Bi). With normalized maximum metabolic rate as the metric, the number of nutrients that gave greater than the overall average rate was counted to show the overlap in utilization between different species (Fig 2Aii and 2Bii). At 37 degrees, 20 nutrients were utilized at above the average rate among all three bacteria tested, while there were groups of nutrients observed to be better utilized in one bacteria alone (31 for *B. cereus*, 15 for *B. anthracis*, and 20 for *S. aureus*). Similar distribution was observed for bacteria incubated in 30 degrees as well, although *B. cereus* once again had the greatest number of nutrients that were utilized (20 for *B. cereus*, 17 for *B. anthracis*, and 14 for *B. subtilis*). As for nutrients metabolized at above the overall average maximum metabolic rates by all bacteria, there were 16 of them at 30 degrees and 20 at 37 degrees. Six of these nutrients were common to both lists (5-keto-D-gluconic acid, D-arabinose, D-ribose, D-xylose, L-arabinose, and L-lyxose) and all of them were either carbohydrates or derivatives (S2 Table). When averages of maximum metabolic rates of nutrients that were well utilized by only one bacteria were compared to that of nutrients well utilized by all bacteria, it was observed that these nutrients resulted in higher rates as compared to nutrients well utilized by one bacteria at 37 degrees (0.68 for *B. cereus*, 0.44 for *B. anthracis*, 0.80 for *S. aureus*, 1.70 for commonly well utilized, p < 0.05, unpaired Student’s t-test) (Fig 2Biii). This may indicate that while choices of carbon utilization are distinct for each species, they also have core parts of metabolism that are common. It is interesting to also note that *B. anthracis* shared more common nutrients with *B. subtilis* at 30 degrees (Fig 2Aii) and *S. aureus* at 37 degrees (Fig 2Bii) than it did with *B. cereus*, which was unexpected. This was also true for *B. anthracis* and *S. aureus* at 37 degrees as compared to *B. cereus*.

**Fig 2.**
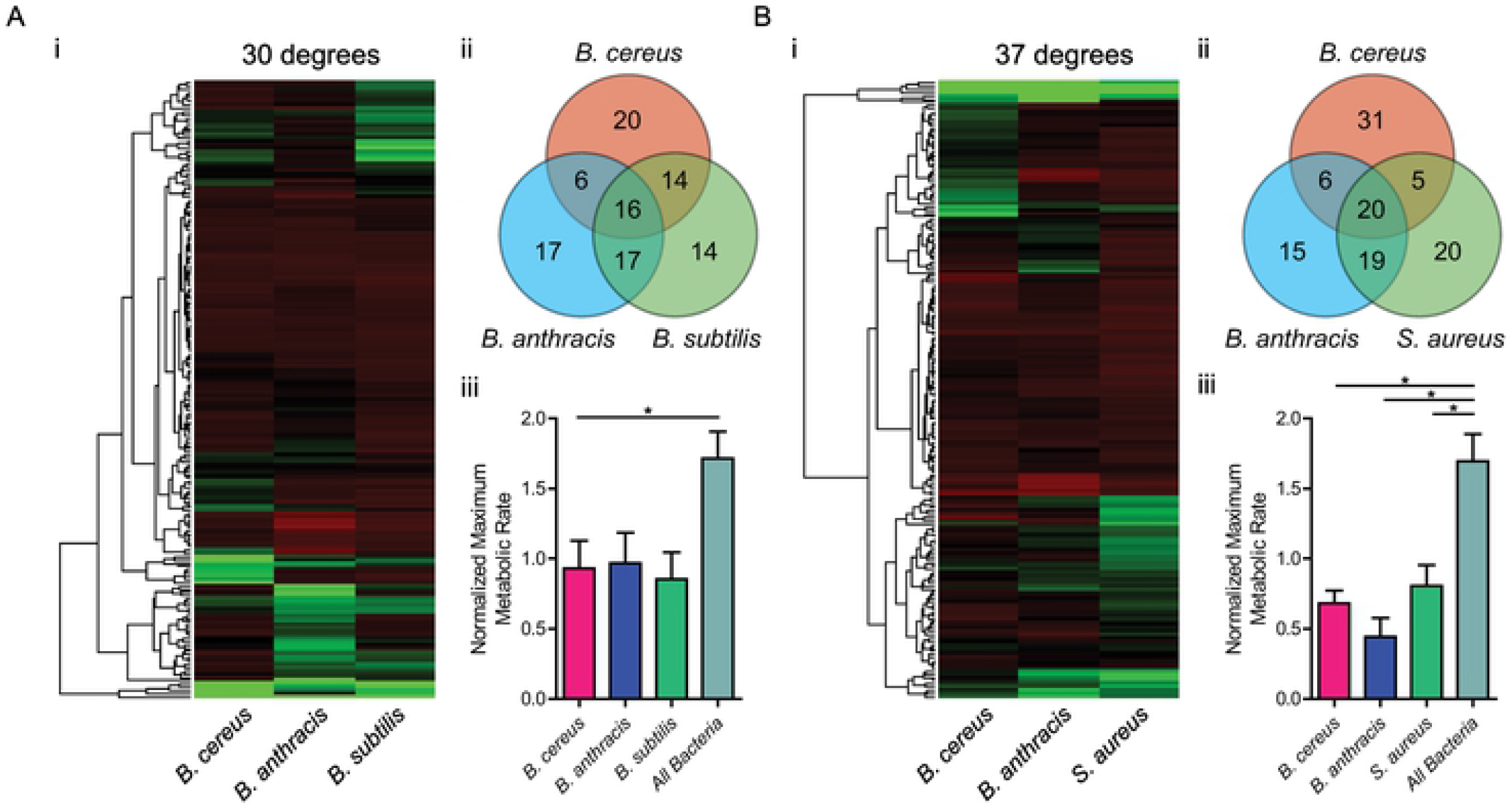
Metabolic rates for carbon sources in bacteria show variations and groupings. (A and B) Maximum metabolic rates of nutrients for bacteria incubated at 30°C (A) and 37°C (B). (i) Nutrients are hierarchically clustered by their chemical structures (dendrograms, left) and metabolic rates observed are shown as heatmaps (right) with each column representing results from different bacteria. (ii) Venn diagrams of nutrients are shown with numbers reflecting the count of nutrients that had metabolic rates statistically greater (p < 0.05) than the overall average rate. Unpaired Student’s t-test was used for comparison. (iii) Normalized maximum metabolic rates for nutrients well utilized by one bacteria are compared against nutrients well utilized by all bacteria. Bars represent averages of all nutrients that had statistically higher metabolic rate than the overall average rate. Error bars represent standard error of the mean. Maximum metabolic rate for each nutrient is an average from three independent experiments (n = 3). *: p < 0.05 by unpaired Student’s t-test.

### Metabolic utilization of nutrients by chemical properties

Nutrients in the plates for carbon metabolism have a wide variety of chemical properties. This fact can be leveraged to determine the types of food bacteria prefer to eat. We classified nutrients into distinct “food groups” based on their chemical properties: carbohydrates, amino acids, lipids, and hydrophobicity according to their calculated partition coefficient (xLogP3) (Fig 3Ai, Bi, Ci, Di) [40]. Every nutrient was queried through NCBI PubChem for assignment into those four criteria and categorized accordingly. Nutrients were hierarchically clustered according to their chemical structural similarities as measured by atom-pair distances using ChemmineR R package within groups [41]. Maximum metabolic rates were standardized to mean of 0 and standard deviation of 1 for each bacteria incubated under their optimal growth temperatures, and visualized as heat maps for comparison, with ‘+’ and ‘-’ indicating groups of nutrients that either belonged or not to the “food group,” respectively (Fig 3Aii, Bii, Cii, Dii). The average maximum metabolic rate for carbohydrates was greater than that of non-carbohydrates for *B. anthracis* (31.32 for carbohydrates, 23.46 for non-carbohydrates, p = 0.0005), *B. subtilis* (16.19 for carbohydrates, 10.80 for non-carbohydrates, p = 0.0030), and *S. aureus* (23.52 for carbohydrates, 16.84 for non-carbohydrates, p = 0.0084, all unpaired Student’s t-test) (Fig 3A). This stands in contrast to amino acids and lipids, where no statistically significant differences were observed between nutrients categorized under these properties (Fig 3B and 3C). As for hydrophobicity, the median value of xLogP for all nutrients, −2.3, was used as the dividing point, with xLogP less than or equal to the median as being deemed relatively hydrophilic and greater as hydrophobic. All four species of bacteria incubated under their optimal temperature had average raw maximum metabolic rates for hydrophilic nutrients greater than hydrophobic nutrients (for *B. anthracis*, ≤ median 29.74 and > median 24.39, p = 0.0183; for *B. cereus*, ≤ median 20.04 and > median 13.53, p = 0.0115; for *B. subtilis*, ≤ median 16.75 and > median 9.67, p < 0.0001; for S. *aureus*, ≤ median 23.28 and > median 16.43, p = 0.0067; unpaired Student’s t-test) (Fig 3D). These results highlight facile metabolic utilization of carbohydrates for these bacteria, as opposed to amino acids and lipids, when bacteria are constrained to primarily one nutrient as their carbon source. Superior utilization of hydrophilic nutrients is also suggestive of carbohydrate metabolism, as 76% of hydrophilic molecules (68 out of 89) are carbohydrates, as opposed to 29% (30 out of 102) for hydrophobic nutrients. Further suggestive of the importance carbohydrates play in carbon metabolism of these bacteria can be observed when chemical formula of the nutrients themselves are examined. When modular arithmetic is applied to the number of carbon atoms in nutrients, there exists statistical correlation between the remainder after divisions by five and six and classification of molecules as carbohydrates when ANOVA is performed (mod 5, p < 0.00072; mod 6, p = 3.74 x 10^−10^). This can be visualized when raw maximum metabolic rates are plotted by their remainders after division by five or six (pentoses have remainder of 0 and 5 after division by 5 and 6, and hexoses have remainder of 1 and 0 after division by 5 and 6), as nutrients that have number of carbon number atoms that fit the modular arithmetic for pentoses and hexoses have greater maximum metabolic rates (mod 5: p < 0.0001; mod 6: p < 0.0001, one-way ANOVA) (S4A and B Fig).

**Fig 3.**
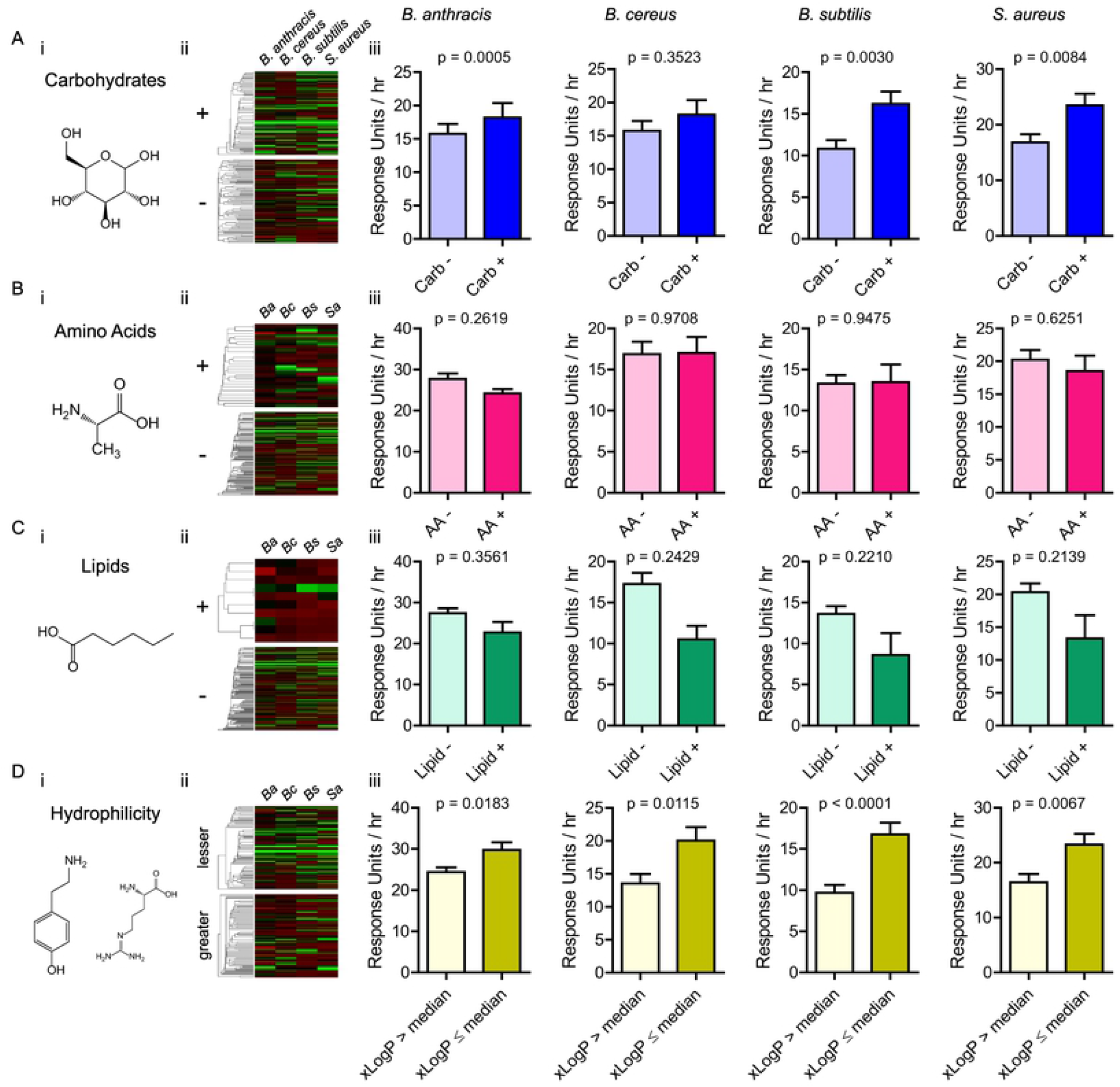
Metabolic rates correspond to certain chemical properties of nutrients. (A-D) Four chemical properties of nutrients examined with the structure of an example from each category (i): (A) carbohydrates (shown: D-glucose), (B) amino acids (shown: L-alanine), (C) lipids (shown: caproic acid), and (D) hydrophilicity as represented by partition coefficient (shown: tyramine and L-arginine). (ii) Heatmaps of maximum metabolic rates for nutrients with nutrients in the category for chemical property under question (+ or lesser) or did not (- or greater). Nutrients are hierarchically clustered by their chemical structural similarities using atom-pair distances. *Ba: B. anthracis, Bc: B. cereus, Bs: B. subtilis, Sa: S. aureus.* (iii) Average maximum metabolic rates for nutrients by chemical property (blue: carbohydrates, red: amino acids, green: lipids, yellow: hydrophilicity / partition coefficient). Bars represent averages of all nutrients categorized by chemical property. Error bars represent standard error of the mean. Maximum metabolic rate for each nutrient is an average from three independent experiments (n = 3). *: p < 0.05 by unpaired Student’s t-test.

### Carbohydrate pathways in carbon metabolism of bacteria

To examine pathways in carbohydrate metabolism for each nutrient, KEGG was queried and each nutrient’s pathway participation was examined [42]. We sought to extract information from metabolic utilization data to determine which metabolic pathways are utilized more efficiently while avoiding *a priori* knowledge of the organism and its metabolic network biasing our analysis. Nutrients classified into pathways involved in carbohydrate metabolism were selected for this analysis, following our determination from the previous section that carbohydrates were preferred than other nutrients. Average standardized metabolic maximum rates for nutrients grouped into 15 different carbohydrate pathways showed overall elevation of utilization across bacteria with the exception of *B. cereus* at its suboptimal temperature of 37 degrees (Fig 4A). The pentose phosphate pathway had the highest normalized maximum rate at 30 degrees for all bacteria (0.917 for *B. cereus*, 1.488 for *B. anthracis*, and 1.239 for *B. subtilis*). Of particular note was the comparison in number of nutrients that showed higher utilization associated with carbohydrate pathways between *B. anthracis, B. cereus*, and *B. subtilis* that emphasize the distinctiveness of *B. anthracis*’ carbohydrate metabolism. At 30 degrees, the top four carbohydrate pathways for *B. anthracis* were amino and nucleotide sugar metabolism, inositol phosphate metabolism, pentose and glucoronate conversion, and pentose phosphate pathway (1, 10, 11, and 12). (Fig 4B). While most of these pathways were found to have high metabolic rates for *B. cereus* and *B. subtilis* as well, an exception was noted for inositol phosphate pathway, which had negative normalized maximum metabolic rate indicating that it was not utilized well by these two bacteria (0.7938 for *B. anthracis*, −0.0464 for *B. cereus*, −0.0605 for *B. subtilis*). At 37 degrees, a similar pattern of carbohydrate pathway utilization was observed for *B. anthracis* and *B. cereus*, with the same pathways being commonly well utilized and inositol phosphate pathway being underutilized in *B. cereus*. For *S. aureus* eleven out of fifteen carbohydrate pathways had positive normalized maximum metabolic rates, hinting at a more diversified use of carbohydrates (Fig 4C). Curiously, the inositol phosphate pathway was not one of the pathways well utilized for *S. aureus* (−0.0583), indicating that its utilization might be specific for *B. anthracis.*

**Fig 4.**
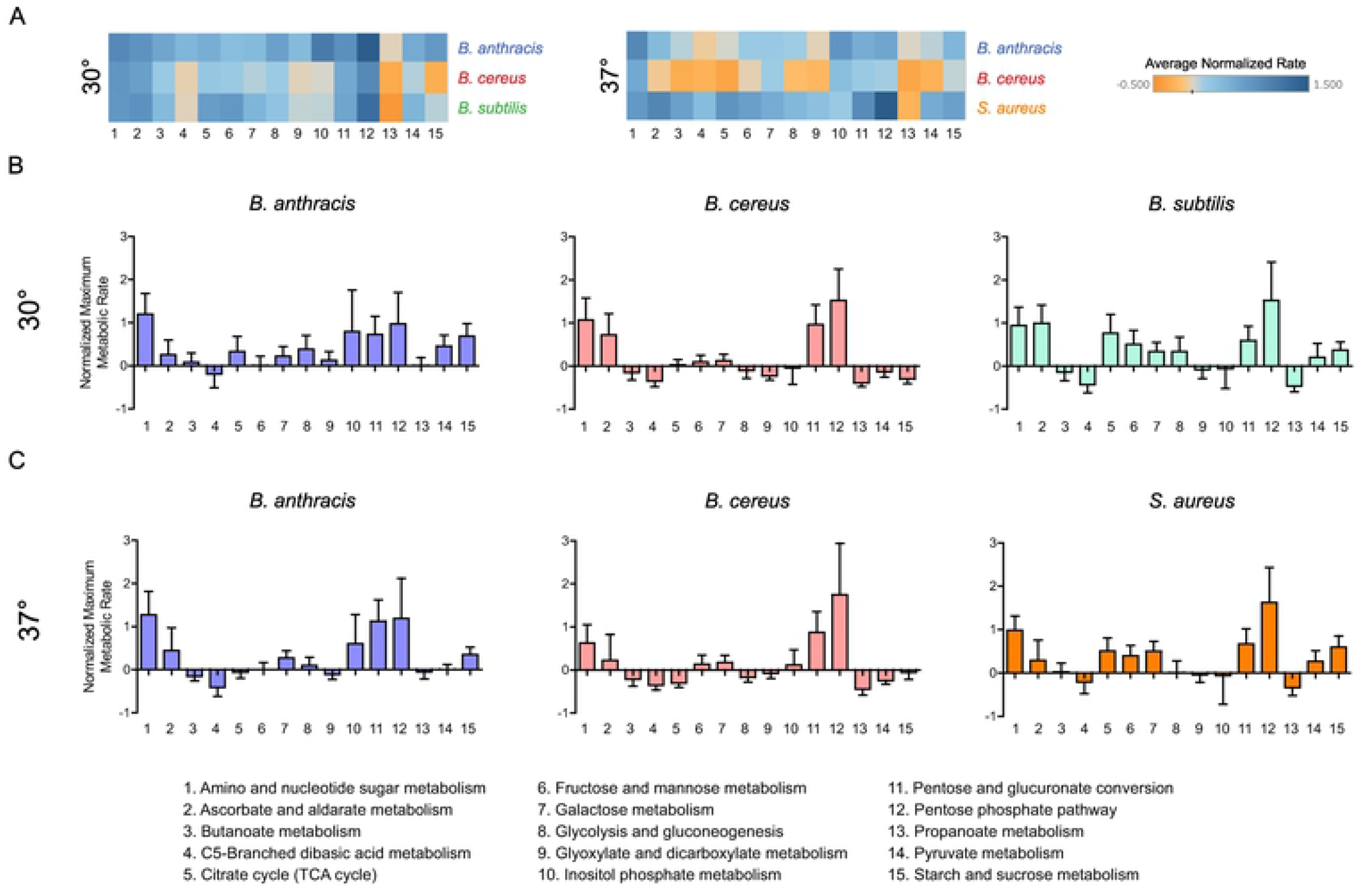
Certain carbohydrate pathways have superior utilization of nutrients. (A) Heatmaps of normalized maximum metabolic rates for nutrients utilized by different carbohydrate pathways. Nutrients are categorized by which carbohydrate pathways they are utilized in, and average of all maximum metabolic rates from nutrients for each carbohydrate pathway are shown as heatmaps. (B and C) Bar graphs of normalized maximum metabolic rates for nutrients in all carbohydrate pathways. (B) shows results from bacteria incubated at 30°C, and (C) shows results from 37°C. Each bar represents average maximum metabolic rates for all nutrients for each carbohydrate pathway. Error bars represent standard error of mean. Maximum metabolic rates are normalized to average of 0 and standard deviation of 1. Each nutrient’s maximum metabolic rate is an average from three independent experiments (n = 3).

### Individual nutrients and their metabolic pathway associations

Analysis of overall averages of metabolic rates suggests that there exist variations in metabolism at the level of individual nutrients. Using metabolic pathway assignments made for every nutrient in the previous analysis, individual nutrients and pathways were ordered by their standardized maximum metabolic rate at 37 degrees and laid out as heat maps for carbohydrate (Fig 5A) and amino acid pathways (Fig 5B). These rates for individual nutrients show that even within pathways that average high metabolic rate for nutrients tested, there exists a large variation of utilization of nutrients within individual pathways (for the pentose phosphate pathway, from 6.279 for 5-keto-D-gluconic acid to −0.663 for D-gluconic acid in *B. cereus*, 4.558 for D-ribose to −0.773 for 2-deoxy-D-ribose for *B. anthracis*, 4.212 for D-ribose to −0.718 for D-glucosaminic acid in *S. aureus* – as examples). There are universally well-utilized nutrients within carbohydrate pathways, such as L-arabinose (4.390 for *B. cereus*, 4.051 for *B. anthracis*, and 2.926 for *S. aureus*), which was consistently involved in the top three out of four pathways (amino sugar and nucleotide metabolism, pentose and glucoronate interconversion, and pentose phosphate pathway). In contrast, analysis of amino acid pathways reflects a more modest degree of utilization and does not show the heterogeneity as observed in carbohydrate pathways. This is more evident when the top and bottom ten nutrients in metabolic maximum rates are separately visualized for carbohydrate metabolism (Fig 5C) and amino acid metabolism (Fig 5D). *B. cereus* and *S. aureus* had a small group of nutrients metabolized exceptionally well even within the top ten (four nutrients with normalized rates greater than 3, which is equivalent to three-fold greater rates than the standard deviation, for *B. cereus –* 5-keto-D-gluconic acid, L-lyxose, D-ribose, and L-arabinose; and two nutrients for *S. aureus* – D-ribose and 5-keto-D-gluconic acid), while *B. anthracis* had seven nutrients with rates that exceeded the threshold rate of 3 (D-ribose, D-glucosamine, D-xylose, L-arabinose, 5-keto-D-gluconic acid, D-arabinose, and L-lyxose). In contrast, none of the nutrients involved in amino acid pathways exceeded the threshold of 3. This high efficiency of metabolism observed for *B. anthracis* in carbohydrate pathways for a larger number of nutrients than *B. cereus* or *S. aureus* suggests that the carbohydrate metabolism of *B. anthracis* would be more efficient in environments with a limited variety of nutrients.

**Fig 5.**
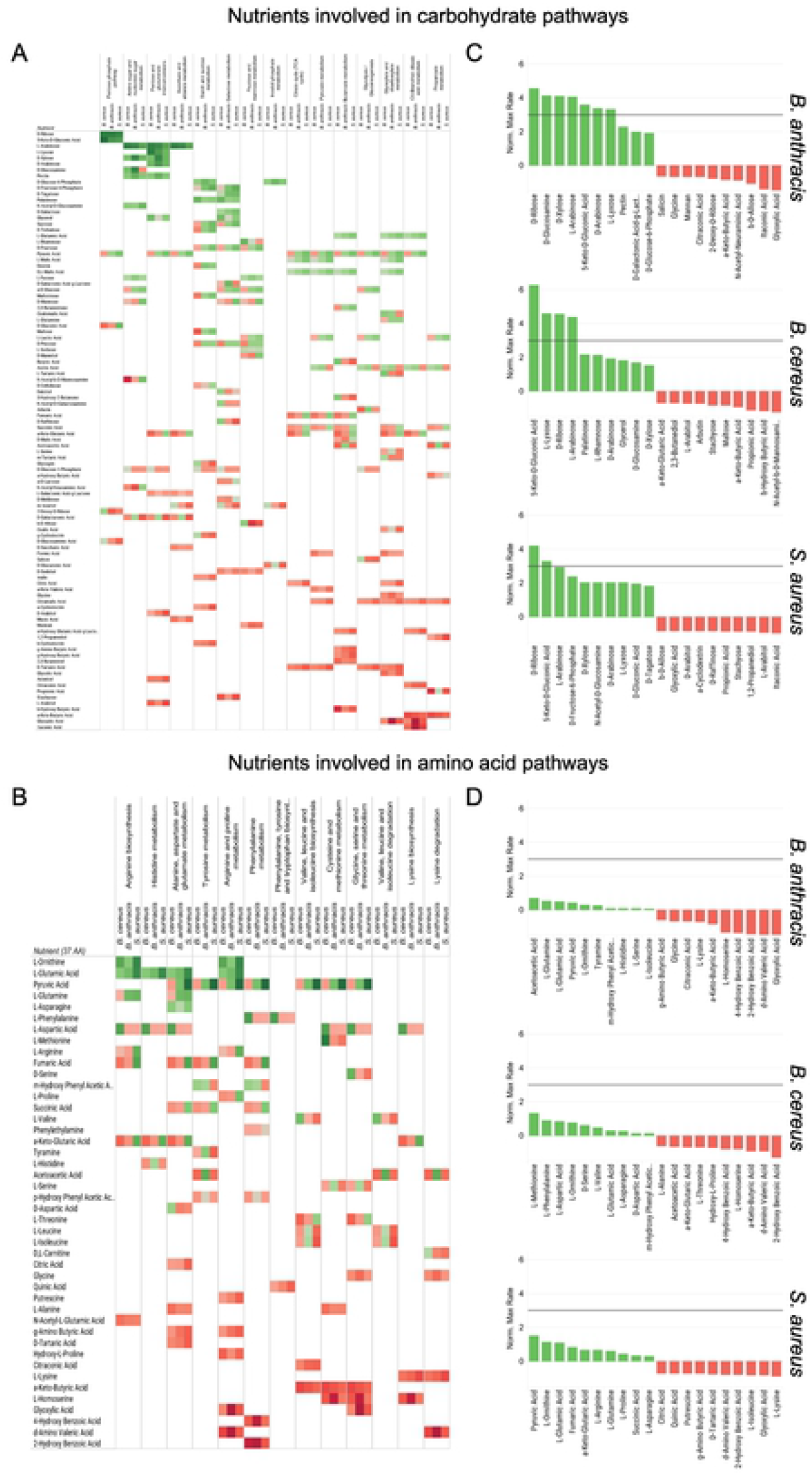
Nutrients are utilized in different pathways with wide range of metabolic rates. (A and B) Heatmap showing normalized maximum metabolic rates for all nutrients associated with carbohydrate pathways (A) and amino acid pathways (B). For every nutrient (left column), normalized maximum metabolic rates for bacteria incubated in 37°C are shown in three columns (*B. cereus, B. anthracis*, and *S. aureus*) for all pathways that the nutrient is associated with. Nutrients are ordered from top to bottom by their overall average metabolic rate. Pathways are ordered from left to right by their average metabolic rate. (C and D) Bar graphs of normalized maximum metabolic rates for nutrients with top and bottom 10 metabolic rates involved in carbohydrate pathways (C) and amino acid pathways (D). For each bacteria, maximum metabolic rates for nutrients with 10 highest metabolic rates are shown in green, and 10 lowest metabolic rates are red. Maximum metabolic rates are normalized to average of 0 and standard deviation of 1. Gray lines indicate normalized rate of threshold of 3, which is equivalent to three standard deviations greater than the mean. Each nutrient’s maximum metabolic rate is an average from three independent experiments. (n = 3).

### Nutritional preferences of *B. anthracis* in serum

To better characterize the global nutrient requirement of pathogenic *Bacillus* under conditions designed to simulate growth in a mammalian host, carbon sources from the screen were supplemented with 40% fetal bovine serum (FBS) and the entire analysis repeated. Nutrients were ordered according to their maximum metabolic rates. Interestingly, nutrients well used in serum by *B. anthracis* were not identical to those in minimal media, with Spearman’s ρ of 0.5050 (Fig 6A). When nutrients are categorized by carbohydrates and amino acids, their utilization essentially flips in serum compared to media (carbohydrates: −0.2474 vs. non-carbohydrates: 0.1809, amino acids: 0.2397 vs. non-amino acids: −0.0700, lipids: −0.0588 vs. non-lipids: −0.0187) (Fig 6B). The most striking observation is that *B. anthracis* no longer utilizes carbohydrates well in serum (or perhaps uses them less), with amino acids now seemingly being the dominant nutrient of choice (carbohydrates: −0.2474, amino acids: 0.2397, p = 0.0264, unpaired Student’s t-test). This change of metabolism is most apparent when comparing the number of nutrients that have higher normalized metabolic rates in media as opposed to those in serum (for carbohydrates: 50 in media vs. 39 in serum, for amino acids: 11 in media vs. 19 in serum). This involvement of pathways in metabolic differences in serum is most readily seen when nutrients themselves are categorized by pathways. Out of 15 carbohydrate pathways catalogued, the average of standardized maximum metabolic rates for nutrients in 8 pathways are negative, whereas 12 out of 13 amino acid utilization pathways average in the positive (Fig 6C). For carbohydrate pathways, average maximum rates range from −0.788 for amino sugar and nucleotide metabolism to 0.661 for citric acid cycle. While lipid metabolism as a category contains both the lowest (−2.540 for fatty acid biosynthesis) and highest rates (1.581 for glycerophospholipid metabolism), no statistically significant trends could be discerned. On the other hand, amino acid metabolism pathways (with the exception of branched chain amino acid degradation), all ranged in positive from 0.106 (lysine degradation) to 0.548 (lysine biosynthesis), reflecting how nutrients involved in amino acid pathways are well utilized. Analyses of these pathways at the nutrient level for carbohydrates (S7A Fig) and amino acid pathways (S7B Fig) show that decreases in metabolic rates for carbohydrates for media to serum are the greatest for certain pentoses (D-xylose: −3.808, D-arabinose: −3.616, D-ribose: −4.243) and hexose derivatives (D-galactonic acid-g-lactone: −2.897, D-glucosamine: −4.053), demonstrating that these simpler carbohydrates, while well utilized in nutrient-poor conditions, no longer become efficient carbon sources for metabolism in environment rich with diverse nutrients. These results suggest that there are massive changes to carbon metabolism that is dependent on the environment the bacteria find themselves in, with *B. anthracis* switching to favor catabolism of amino acids in serum.

**Fig 6.**
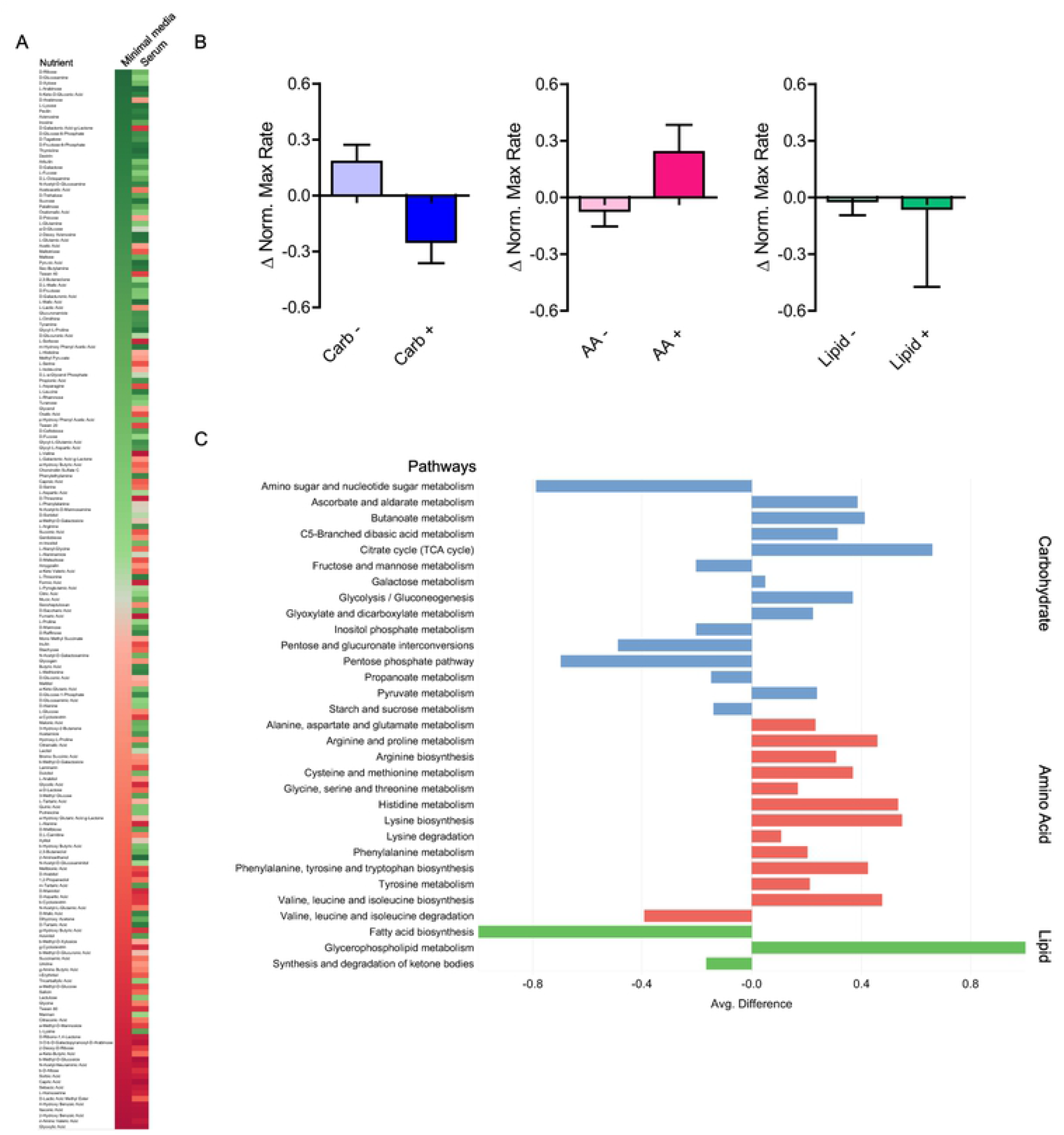
*B. anthracis* has metabolic profile dependent on nutrient availability. (A) Comparison of ordered lists of maximum metabolic rates between nutrient restricted (minimal media) and enriched (serum) environments. Color gradient shows rank of nutrients by their metabolic rate. The ordered list from nutrient restricted condition (left) is shown ordered, and corresponding rank from nutrient enriched condition (right) is placed side as comparison. (B) Differences of average metabolic rates between nutrient-restricted and enriched conditions by nutrient category. Bar graphs show differences between average maximum metabolic rates for nutrients by their categorization (blue: carbohydrates, red: amino acids, green: lipids). Error bars represent standard error of the mean. (C) Differences of average metabolic rates by pathways associated with nutrients. For each pathway, differences in maximum metabolic rates of all nutrients associated with that pathway between nutrient restricted and enriched conditions were averaged and shown as a bar graph. Colors represent pathway categories (blue: carbohydrates, red: amino acids, green: lipids).

## Discussion

From this study, we are able to establish that: i) metabolic activity of bacteria can be measured colorimetrically through chemical reduction potential, ii) nutrients have different degrees of utilization among different bacteria, iii) the choice of which nutrients to use is impacted by temperature; generally, the nutrient preferences track with whether the species grows in the environment versus the host, iv) the chemical properties of the nutrients affect their metabolic utilization rate; carbohydrates and hydrophilic nutrients are generally preferred in media, v) within carbohydrate metabolism, higher metabolic rates are limited to few specific pathways that use a handful of same nutrients, and vi) in serum, *B. anthracis’* nutrient preferences are vastly different then in defined media; mainly, the nutrient preference shifts from carbohydrates to amino acids.

Infection of a host by bacteria requires these pathogens to be adaptable metabolically in nutritionally austere environments. One component of a host’s nutritional immunity to starve out pathogens would be to keep down the level of free amino acids and lipids. Pathogens would then be expected to tune their metabolism to use freely available nutrients such as carbohydrates. Since previous studies have shown that pathogens thrive in carbohydrate-rich environments, we expected to observe a high degree of metabolic utilization for carbohydrates in general; our data reinforces this notion [43-46]. However, the type of carbohydrate each species preferred varied substantially. The data suggests that not all carbohydrate metabolic pathways are equally tuned for utilization. Indeed, there would be resource costs involved in creating and maintaining metabolic pathways that remain unused or underutilized, and these pathogens would only need to have in preparation pathways involved in metabolizing nutrients frequently encountered during their lifecycle. For the experiment performed in auxotrophic media, where a single nutrient is the predominant source of carbon, bacterial maximum metabolic rate measured reflects the readiness of bacteria’s metabolic pathways to utilize that nutrient. This raises a point noted during both nutrient and pathway analyses: why are pentoses utilized better than other forms of carbohydrates in *B. anthracis*? High metabolic utilization observed for nutrients involved in the pentose phosphate pathway offers an explanation. In addition to being catabolized for energy production, these pentoses can also be readily used for anabolism to build up metabolic machinery through the pentose phosphate pathway. These newly synthesized metabolic components in turn allow for even better utilization of pentoses provided in the environment, creating a positive feedback loop that allows *B. anthracis* to thrive in a nutrient limited environment. *B. anthracis* would normally encounter nutritionally restricted surroundings during parts of its infectious cycle, such as attempting to survive within a macrophage’s endosomes. *B. anthracis* relies on toxins to further progress in its course of infection, and ability to remain metabolically active in nutritionally deficient condition would be valuable. Nutrients that result in high metabolic yield under an ideal condition may not be the best nutrient for every situation, especially when bacteria must deal with resource-poor environment. This may be the reason why certain hexoses such as glucose, which are previously known to be well utilized by various pathogens during infection, do not result in particularly high metabolic rate when they are provisioned as the sole source of carbon. Conversely, in the nutrient-rich environment tested in this experiment with 40% fetal bovine serum, bacteria no longer need to focus on taking a balanced approach; instead, maximum metabolic rate is primarily determined by total capacity for metabolism. In fetal bovine serum supplemented media, *B. anthracis* is no longer restricted to one carbon source for both energy and anabolism, and the nutrient screen thus serves as a proxy for which nutrients allow for most expansion of metabolic pathway capacity. This would result in nutrients utilized in amino acid metabolism generally giving higher metabolic rates, as these nutrients would be directly used in anabolism to expand metabolic pathways to allow higher maximum rates. This hypothesis will need to be tested.

Our results show that the chemical properties of the nutrient can correlate with their metabolic utilization. One intriguing chemical property we observed is the partition coefficient for nutrients (LogP), which is a quantitative measure of a physical property of molecules as opposed to a descriptive categorical variable. Many cases of high metabolic rates observed for low partition coefficient molecules could be explained by the presence of well-metabolized carbohydrates, which are hydrophilic molecules. However, there were exceptions to this as seen by nutrients with low partition coefficients that are carbohydrate-derivatives but still not utilized well by bacteria. On the other hand, hydrophobic molecules with their larger number of energetic carbon-carbon bonds may initially appear as energy-dense molecules that yield higher metabolic payoff. However, considering the fact that pathogenic *Bacillus* must spend majority of its lifecycle inside the animal host with water as the primary matrix, it follows that these *Bacillus* would be proficient at uptaking and utilizing nutrients that are hydrophilic and water soluble (with low partition coefficient). High metabolic utilization for bacteria implies that a transport mechanism already is in place to import these nutrients from the environment, as well as having pathogens ready to convert the primary metabolism’s products into biological building blocks for anabolism. Given that these *Bacillus* are much more likely to encounter these hydrophilic nutrients during a course of infection, they would also have ways to utilize these nutrients. Conversely, these hosts also employee nutritional immunity during bacterial infection to counter these *Bacillus* and other bacterial pathogens. Studies have demonstrated that abundance of glucose during bacterial infection correlate with poorer outcome, while switching of the host’s metabolism away from carbohydrates towards lipid and amino acid consumption can aid the host in battling the infection [43-45, 47-50]. This role nutrition plays during infection has been principally investigated in context of host immunity and inflammation, but our study suggests that this topic also merits consideration from the perspective of bacterial nutrition consumption as well [51-54]. Our findings indicate that through the readiness of *Bacillus* for such nutrients, partition coefficient of nutrients is another one of factors that can influence growth of a pathogen in the host. We demonstrate in this study that approaching nutrients as a limited resource that must be utilized by pathogens offers another perspective to host-pathogen interaction, especially in the context of all the nutrients that are available at that time.

The connection between nutrition and bacterial infection has been so far primarily approached from the perspective of host malnutrition and dysfunction of host’s immune response, as it had been assumed that bacteria are indiscriminate in their preferences to utilize all possible categories of nutrients [55]. And while the host’s responses regarding nutrients occur at the organismal level, direct nutritional tug-of-war between host and bacterial pathogens occurs at the molecular level. As our study demonstrates, bacterial pathogens are more metabolically proficient when consuming certain nutrients and it is reasonable to expect them to be more pathogenic toward the host when encountering an optimal combination of nutrients. One well-characterized component of host’s nutritional immunity is the sequestration of key micronutrients, such as iron, which removes these linchpins of metabolism from being accessible to pathogens through biochemical means [5,6]. Is it possible that mammalian hosts deploy a similar strategy with macronutrients? In humans, the concentrations of various amino acids in blood are kept in the micromolar range. Given the heightened metabolic utilization of amino acids and associated nutrients by *B. anthracis* in serum observed here, keeping the concentration of amino acids low could also be a part of nutritional immunity. On other hand, physiological concentrations of carbohydrates can be in the millimolar range. While situationally lowering this already high concentration of carbohydrate in blood as a part of response against infection might be impractical for the host, more achievable would be to lower the amino acid concentrations, which may adversely affect pathogen protein anabolism. In this study, we also demonstrate that even among the same class of nutrients, metabolic utilization can vastly differ from one nutrient to another. This suggests that when either depriving or interfering with a bacterial pathogen’s metabolism of nutrients, only targeting a handful of nutrients and pathways that have high utilization might be sufficient for disarmament. Unlike antibiotics that target one component of a bacterial cellular process (usually essential), this method of metabolism control would aim to shut down a bacterium’s ability to derive energy or build larger biomolecules.

As pertaining to the direct control of metabolism, a handful of nutrients were metabolized with maximum rates that were much lower than the average of all nutrients screened. While some nutrients by definition were expected to be metabolized at lower rates than the average, observations that some of these nutrients also had equally lower metabolic maximum rate in the enriched condition as tested with serum was surprising, for this indicated that metabolism of *B. anthracis* was outright slow with these nutrients. There are two possible explanations for this decreased metabolic activity in the presence of these nutrients: one is that nutrients themselves are utilized at a slower rate, and decreased maximum rate observed is due to lack of additive effects normally found between the nutrients and enriched media. More intriguing possibility is that nutrients directly interfere with metabolic consumption of other resources from enriched media. Nutrients involved in amino acid metabolism resulted in faster metabolic maximum rates than the overall average for *B. anthracis* (Fig 6C, S7B and S7C Fig), and only a limited number of nutrients (21) had higher rate than the negative control without supplemental nutrients. Given these two facts, it stands to reason that *B. anthracis* in a nutritionally plentiful environment can be selective as to which nutrients to utilize in metabolism and choose to leave alone nutrients that would not result in efficient usage. While this may explain why the majority of nutrients supplemented did not result in increased maximum rate, there were five nutrients where the maximum rate did not even reach 30% of the negative control: capric acid, β-methyl-D-glucoside, glyoxylic acid, 2-hydroxy benzoic acid, and itaconic acid. This decrease in the maximum metabolic rates supports the scenario where the interference of metabolism by the nutrient itself must be considered as a possible cause for this decrease in the metabolic maximum rate. How could the antagonistic relation between these added nutrients and enriched pool of chemical resources occur? For these five nutrients, not one classification of pathways seems to explain the reason why these degrade metabolism. Glyoxylic acid features prominently in multiple pathways as a key component of glyoxylate shunt, an alternative pathway to citric acid cycle, but other four nutrients are not widely utilized at all [56]. This dichotomy in pathway utilization hints that there could be two distinct ways in which these nutrients adversely affect the metabolism. One possible explanation is that the nutrient itself directly acts as an inhibitor of metabolic enzymes. Given that these four nutrients are not normally used as metabolites, it can be argued that these molecules may be inhibitors that slow down metabolism as either drug-like or signal molecules. Indeed, in case of itaconic acid, the ability of such chemical derivatives of metabolites to inhibit the bacterial growth through metabolic interference has been previously demonstrated [57]. The other explanation, which may be the reason for glyoxylic acid, is that these nutrients themselves tune down the metabolism through feedback. In case of glyoxylic acid, a key piece in glyoxylate shunt which is operated to synthesize carbohydrates from other carbon sources when *B. anthracis* is only provisioned with non-carbohydrate carbons, proper functioning of overall metabolism might not be possible even when the pathway itself is present [58,59]. While it remains to be seen whether it would be feasible to achieve pharmacologically relevant local concentrations of these metabolism antagonists at the bacterial level to block the proliferation of *B. anthracis in vivo*, our study offers glimpses into how such strategy could be utilized when these nutrients are applied as antibiotics.

At the level of organism, chemical categories of nutrients do not seem to be specific enough to distinguish one bacteria from another by metabolic performance alone. Rather, it is at the pathway level where nutrient utilization can differentiate one bacteria to another. While most nutrients were either universally well-utilized or poorly-utilized, there still were a number of nutrients where high utilization was restricted to one species. These species-level utilization signatures were found across multiple nutrient utilization pathways, indicating that their uniqueness arose as a part of bacteria’s specialization into their ecological niches with specific mixture of nutrients. While *B. anthracis* spends only a part of its lifecycle in soil, one which is considered metabolically inert (the spore), it is widely accepted that *B. cereus* and *B. subtilis* thrive on soil [13,17,20]. All three bacteria share soil as the backdrop for a big part of their lifecycle, yet they still differ greatly in metabolic utilization of same nutrients. This hints that their metabolic specializations arose not just as a product of nutrient availability, but their interactions with host organisms as pathogens as well. This raises the intriguing possibility of tailoring therapeutics using nutrient classes that can specifically target metabolic specializations at a species-specific level. This might be especially useful at selectively targeting certain pathobionts amongst a myriad of beneficial or non-pathogenic commensal species, for example, in the gastrointestinal tract.

## Materials and methods

### Preparation of bacteria for assays

Frozen bacterial stocks of *B. anthracis* Sterne, *B. cereus* 10987 (ATCC), *B. subtilis* 2091 (ATCC), and *S. aureus LAC* were added to 1 mL of Luria Bertani (LB) media at 1% inoculum and incubated in 30°C or 37°C overnight with 160 rpm orbital shaking to stationary growth phase (OD_600_ > 1.5). Kanamycin was added to the medium for selection (50 μg/mL for *B. anthracis* and *B. cereus*). One mL of bacterial culture was washed twice with 1 mL of deionized water after spinning down in Beckman Coulter centrifuge (Indianapolis, IN, USA) for 3 minutes at 17000xG. Washed cells were diluted in IF-0a inoculating fluid from Biolog (Hayward, CA, USA) – here referred to as minimal media – to 81% transmittance equivalent (OD_600_ ∼0.093) as measured by spectrophotometer (Beckman Coulter).

### Bacterial growth assay

For growth on 96-well Phenotype MicroArray™ carbon utilization assay plates (Biolog), 880 μL of washed bacterial cells were added to the assay media of following composition: 10 mL of IF-0a inoculating fluid (Biolog) and 1.12 mL of deionized water for total volume of 12 mL. The list of nutrients in Phenotype MicroArray™ carbon utilization assay plates can be found in supplemental information (S1 Table). Well number 7 of the Phenotype MicroArray plate 2 contained gelatin, which due to its heterogenous composition was excluded from all further analysis. The assay media had following concentration of additives: 2 mM MgCl_2_·6H_2_O, 1 mM CaCl_2_·2H_2_O, 25 μM L-arginine HCl, 50 μM L-glutamine Na, 12.5 μM L-cystine, 25 μM 5’-UMP 2Na, 0.005% yeast extract, and 0.005% Tween 80. 100 μL of bacterial cells in the assay media were dispensed into each well of Phenotype MicroArray™ plate, and plates were incubated at stationary position in 30°C or 37°C for 24 hours in Synergy™ plate reader (BioTek, Winooski, VT, USA) with 550 and 600 nm absorbance readings taken every 15 minutes.

### Metabolic utilization assay

To measure metabolic utilization of various carbon sources by bacteria, Phenotype MicroArray carbon utilization assay plates were prepared in the same fashion as in the growth assay, but also with 120 μL of tetrazolium-based dye mix F (Biolog) added to the assay media to total volume of 12 mL. 100 μL of bacterial cells in the assay media were dispensed into each well as previously. Plates were incubated in OmniLog™ plate reader (Biolog) in static position at 30°C or 37°C for 24 hours, with metabolic activity reflected by the color change of dye from transparent to purple. Measurements of color changes were made every 15 minutes. Resulting raw data was first aggregated and processed through OmniLog PM™ program (Biolog) and exported as comma-separated values files for further analysis.

### Extraction of metabolic endpoints and rates

Raw metabolic data in form of comma-separated values file was imported into MATLAB (Mathworks, Natick, MA, USA) to obtain metabolic endpoints and maximum metabolic rates for each nutrient. Metabolic endpoint was defined as the increase of the metabolic value from the value of the metabolic curve at the beginning of its increase in value throughout the course of experiment and exponential moving average (EMA) of the metabolic curve at the conclusion of experiment. The threshold value for the beginning of metabolic curve increase was defined as metabolic activity at the timepoint when the metabolic value was 10% greater than the average of all previous timepoints. EMA was determined by the formula:

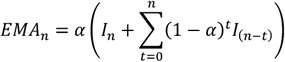

where I is the raw intensity reading, n is the number of datapoints, and α is the weighing coefficient which was set as 0.25 [60].

To determine the maximum metabolic rate from the curve of color change over time, the rate of metabolism value change over the entire experiment was calculated and the largest rate change defined as the maximum metabolic rate. Fifth degree polynomials were fitted to raw metabolic curves using the MATLAB function polyfit to minimize the error from stochastic variations in metabolic curves from one time point to next. The polynomial generated by curve fitting was differentiated with diff function to symbolically derive a function of metabolic rates, a table of metabolic rates at all time points generated, and the maximum value from the metabolic rate table chosen.

### Hierarchical clustering of nutrients

For hierarchical clustering of by the chemical structure, chemical structures for nutrients in Phenotype MicroArray™ carbon utilization screen were queried from PubChem Download Service as SDF files. SDF files were converted to atom distance pairs using R v3.5.3 with the package ChemmineR’s sdf2ap function, and fpSim function was used to calculate similarities and generate a distance matrix. The distance matrix of chemical structural similarities was used for R’s hierarchical clustering function hclust and visualized with heatmap.2. For hierarchical clustering by metabolic data, metabolic data was directly used to calculate a set of pairwise distances by MATLAB function pdist. Euclidean distance was used as the distance metric. Pairwise distances between nutrients were converted into a square matrix with function squareform. Resulting distance matrix generated was clustered with R function hclust and visualized with the function heatmap.2 [61].

### Fetal bovine serum-supplementation metabolic assay

880 μL of washed bacteria suspended in IF-0a media were added to 4.8 mL of fetal bovine serum (Gibco), 120 μL of dye mix F, and 6.2 mL of phosphate buffered saline, pH 7.8, for the final fetal bovine serum concentration of 40% v/v. 100 μL of this bacterial suspension in 40% fetal bovine serum was added to each well of Phenotype MicroArray™ carbon utilization plates, and plates were incubated at static position in 30°C or 37°C for 24 hours in Synergy™ plate reader (BioTek) with the color change due to metabolic activity measured as 550 nm absorbance readings taken every 15 minutes. To verify that 550 nm absorbance reading correlated with metabolic activity obtained in the metabolic utilization assay, raw data from the Phenotype MicroArray™ plate 2 from both metabolic utilization experiments were plotted linearly and R^2^ value calculated to confirm the degree of correlation (Data not shown).

### Statistical analysis

Unpaired Student’s t-test and one-way ANOVA with Tukey post-hoc test were performed on GraphPad Prism (GraphPad Software, La Jolla, CA, USA). Spearman’s rank correlation coefficients were calculated with Excel. Principal component analysis of metabolic data was performed with R’s prcomp function and visualized with fviz_pca_ind. On all statistical analysis, P-values less than or equal to 0.05 were considered significant and marked with an asterisk in the graphs. All visualization was performed through GraphPad Prism, R, or Tableau (Tableau Software, Mountain View, CA, USA).

## Acknowledgements

The authors thank Dr. Zachary Conley for his assistance on data acquisition. The authors thank Drs. Nina Poole and Justin Clark for valuable discussion and feedback. This work was supported by grants AI125778, AI133001, and AI097167 from the National Institute of Health, Allergy and Infectious Diseases Division. The authors declare no conflicts of interest.

## Supporting Information

**S1 Fig. Maximum metabolic rates of bacteria in all conditions for all nutrients**

The full list of all nutrients examined in this study is shown with maximum metabolic rates for all bacteria and conditions tested. Darker shades reflect higher rates, and lighter shades lower rates.

**S2 Fig. Comparing rank lists for maximum metabolic rates and growth as measured by OD**_**600**_ **for three *Bacillus* species**

For *B. anthracis, B. cereus*, and *B. subtilis*, corresponding rank lists for maximum metabolic rate (left) and OD_600_ (right) are shown. Darker tones show higher ranking with higher metabolic rate and OD_600_, and lighter tones show lower ranking.

**S3 Fig. Maximum metabolic rates and metabolic endpoints for all nutrients**

For all bacteria and incubation temperatures (37°C: red, 30°C: blue) investigated in this study, maximum metabolic rate (A) and metabolic endpoints (B) observed for all nutrients are shown as box and whisker plots. Each dot represents an average of metabolic data observed for one nutrient. Whiskers represent 5^th^ and 95^th^ percentile range, while boxes represent 25^th^ and 75^th^ percentile with the middle line representing the median. Each nutrient’s metabolic data is an average from three independent experiments (n = 3).

**S4 Fig. Maximum metabolic rates by modular arithmetic on number of carbon atoms in nutrients**

Maximum metabolic rates for nutrients are grouped by remainders after dividing number of carbon atoms in the nutrient by 5 (A) or 6 (B). Data shown are combined from bacteria incubated in their optimal temperature (37°C for *B. anthracis* and *S. aureus*, 30°C for *B. cereus* and *B. subtilis*). Each nutrient’s metabolic data is an average from three independent experiments (n = 3). One-way ANOVA was performed for p-values and Tukey’s range test was used for pairwise comparisons (*: p < 0.05, **: p < 0.005, ***: p < 0.001).

**S5 Fig. Role of temperature in maximum metabolic rates observed by nutrient property**

Average maximum metabolic rates for nutrients by category are shown as bar graphs. Nutrient properties examined are carbohydrates (A), amino acids (B), lipids (C), and hydrophilicity / partition coefficient (D). Lighter shades represent average rates from 30°C, and darker shades from 37°C. Error bars represent standard error of the mean. Maximum metabolic rate for each nutrient is an average from three independent experiments (n = 3). p-values were obtained with unpaired Student’s t-test.

**S6 Fig. Maximum metabolic rates for nutrients by pathways from bacteria incubated at 30°C**

(A and C) Heatmaps showing normalized maximum metabolic rates for all nutrients associated with carbohydrate pathways (A) and amino acid pathways (C). For every nutrient (left column), normalized maximum metabolic rates for bacteria incubated in 30°C are shown in three columns (*B. cereus, B. anthracis*, and *B. subtilis*) for all pathways that nutrient is associated with. Nutrients are ordered from top to bottom by their average metabolic rate. Pathways are ordered from left to right by their average metabolic rate. (B and D) Bar graph of normalized maximum metabolic rates for nutrients with top and bottom 10 metabolic rates involved in carbohydrate pathways (B) and amino acid pathways (D). For every bacteria, maximum metabolic rates for nutrients with 10 highest metabolic rates are shown in green, and 10 lowest metabolic rates are shown in red. Maximum metabolic rates are normalized to average of 0 and standard deviation of 1. Each nutrient’s maximum metabolic rate is an average from three independent experiments (n = 3).

**S7 Fig. Differences in *B. anthracis* metabolic profile between nutrient restricted and enriched environments**

(A and B) Maximum metabolic rates of nutrients associated with carbohydrate pathways (A) and amino acid pathways (B) are shown as heatmaps. Rates from nutrient restricted environment (minimal media, left), nutrient enriched environment (serum, middle), and difference between two (right) are shown. Nutrients associated with more than one pathway are listed in all associated pathways.

**S1 Table. List of nutrients in Phenotype MicroArray carbon utilization screen**

**S2 Table. List of nutrients with normalized maximum metabolic rate greater than zero for all bacteria**

